# Nuclear and mitochondrial genome assemblies of *Indrella ampulla*, a terrestrial gastropod endemic to the Western Ghats

**DOI:** 10.1101/2025.07.14.664635

**Authors:** Gopi Krishnan, Maitreya Sil, N. A. Aravind, Govindhaswamy Umapathy, Aniruddha Datta-Roy

## Abstract

*Indrella ampulla* is a polymorphic, brightly coloured, forest-dwelling land snail endemic to the Western Ghats of India. Studying and understanding evolutionary processes occurring within this species has remained a challenge largely due to a paucity of genomic resources. We present high-quality annotated nuclear and mitochondrial genome assemblies of *I. ampulla*. The nuclear genome is assembled through a hybrid approach using Illumina short reads and Oxford Nanopore long reads, with an N50 value of 632 kb and 90.3% BUSCO genome completeness. The mitogenome is 13,887 bp long. The demographic history reconstruction based on the genomic data exhibits signatures of population decline during the last 100,000 years. This genome will aid in deciphering the colour polymorphism in *I. ampulla* and augmenting the general understanding of the evolution of gastropods.

**Significance:** *Indrella ampulla* is a terrestrial gastropod endemic to the Western Ghats biodiversity hotspot, and exhibits color polymorphism. The genome of *I. ampulla* provides a valuable resource for understanding molluscan biology, coloration genetics, adaptation, and other evolutionary processes. Our historical demographic analysis offers insights into past population dynamics of this species endemic to the Western Ghats of India, contributing to broader research on biodiversity patterns in tropical ecosystems. The bioinformatics pipeline used in this study provides a reliable framework for assembling high-quality genomes with long-read coverage as low as 10X, making it especially useful for gastropods and other species where obtaining high coverage data is a challenge. Overall, this genome will complement future studies in phylogenomics, comparative genomics, and conservation of molluscs.

## Introduction

Gastropod molluscs are important model systems in evolutionary biology and are the only molluscan group that occupies a wide range of habitats, including marine, estuarine, freshwater, and terrestrial environments (Hayes et al. 2009; Ponder et al. 2019). They exhibit unique traits such as torsion (where internal organs twist along an axis), the presence of toxins, polychromatism in both their shells and their bodies, and a coiled shell – a feature that has been reduced or lost in certain lineages. These adaptations, along with diverse feeding strategies, including herbivory, predation, and parasitism, make gastropods an ecologically and evolutionarily significant group.

Despite their considerable diversity (Aravind et al. 2005), genomic resources for gastropods, particularly those from South Asia, remain limited. This lack of genomic resources hinders the ability to explore key evolutionary and ecological questions within this group. Furthermore, invasive species pose a significant threat to native biodiversity and gastropods are one of the most prominent invasives worldwide. To address this gap, we present the first high-quality hybrid *de novo* genome assembly of a monotypic Indian terrestrial gastropod species, *Indrella ampulla*, a large, semi-slug endemic to the central and southern Western Ghats mountain range (Surya Narayanan and Aravind 2022; Chakraborty et al. 2024).

The species, *I. ampulla* exhibits three brightly coloured morphs (orange, red and yellow), a trait of significant biological interest (Chakraborty et al. 2024). Colouration plays a crucial role in the animal kingdom, serving various ecological and evolutionary functions. While cryptic colouration helps animals evade predation by blending into their surroundings, bright colouration often signals sexual maturity and plays a key role in sexual selection. Additionally, it can function as a warning signal indicating unpalatability or toxicity (Cuthill et al. 2017). The genetic basis of complex traits such as colouration often involves multiple, sometimes previously unknown genes (San-Jose and Roulin 2017).

Here, we report high-quality nuclear and mitochondrial genome assemblies of *I. ampulla* using a hybrid approach that integrates Illumina short and Oxford Nanopore long reads. This genome provides a foundational resource for studying gastropod evolution, colouration genetics, and adaptation in terrestrial environments.

## Methods

### 1. Sample collection and sequencing

The focal species, *Indrella ampulla* is distributed in the Southern and Central Western Ghats which spans across parts of Karnataka, Tamil Nadu, and Kerala states in Southern India (Surya Narayanan and Aravind 2022). For genome assembly, we selected an individual belonging to the red colour morph. The specimen was collected from a cardamom plantation in Nedumkandam, Kerala, and preserved in absolute ethanol. Genomic DNA was extracted from the foot muscle tissue using a modified CTAB extraction protocol (Chakraborty et al. 2020). DNA concentration was assessed both visually on an agarose gel and quantitatively using a Qubit fluorometer. To generate short-read data, libraries were prepared using Illumina’s TruSeq DNA PCR-Free kit, and sequencing was performed on an Illumina NovaSeq 6000 device at Novelgene Inc., Hyderabad. Libraries for long-read sequencing were prepared using Oxford Nanopore Technology’s Ligation Sequencing Kit V14 (SQK-LSK114), and sequencing was carried out on a PromethION device at CSIR-Centre for Cellular and Molecular Biology, Hyderabad.

### 2. Nuclear genome assembly

We generated 21.3 Gb of long-read data (3,393,199 reads) and 136 Gb of short-read data (446 million paired-end reads). Short reads were quality-trimmed using fastp v0.23.2 (Chen et al. 2023), while the long reads were base-called with Guppy Basecaller and error-corrected using the trimmed short reads with fmlrc2 (Mak et al. 2023). Both corrected long-read and short-read datasets were then used for genome assembly. First, we performed a long-read-only assembly using Flye v2.9.4-b1799 (Kolmogorov et al. 2019), resulting in Assembly 1. Next, we generated a hybrid genome assembly incorporating both short and long-read data using MaSuRCA v 4.1.0 (Zimin et al. 2013), producing Assembly 2. Both preliminary assemblies were further polished using the corrected short-read data with Polca pipeline of MaSuRCA. To improve contiguity, we performed two rounds of QuickMerge v0.3 (Chakraborty et al. 2016), combining Assemblies 1 and 2 in different configurations. In the first round, each assembly was used as both reference and query in separate runs. The resulting assemblies were then merged again in a second round of QuickMerge. The specific assembly combinations and their metrics are detailed in Supplementary table 1.

Assembly quality was assessed using Quast v5.2.0 (Gurevich et al. 2013), and completeness was evaluated with BUSCO v5.7.1 (Manni et al. 2021), against the Mollusca database (mollusca_odb10). The final assembly was selected based on an optimal balance of BUSCO completeness and contiguity. The assembly was screened for contamination using NCBI Foreign Contamination Screen (FCS), and the identified prokaryotic sequences were subsequently removed. The assembly summary was visualised using BlobToolKit v4.4.0 (Challis et al. 2020).

### 3. Repeat masking and genome annotation

We first generated a *de novo* repeat library for the genome using RepeatModeler v2.0.5 (Flynn et al. 2020). The genome was then soft-masked with this custom library using RepeatMasker v4.1.4 (Smit et al. 2013) in conjunction with the LTRstruct pipeline. For gene prediction, we used GALBA v1.0.11 (Bruna et al. 2023), incorporating 1.18 million molluscan protein sequences from the UniProt database as external evidence. The resulting annotation was refined by removing unsupported genes and retaining only the longest isoforms. Proteome completeness was assessed using BUSCO in protein mode against the Mollusca dataset, while the proteome quality was evaluated with OMArk v0.3.0 (Nevers et al. 2024) against the Luca.h5 dataset. Functional annotation of the proteome was performed using EggNOG-mapper (emapper v2.1.12; Cantalapiedra et al. 2021), utilising EggNOG orthology data (eggNOG 5.0; Huerta-Cepas et al. 2019) for comprehensive annotation.

### 4. Mitogenome assembly

We assembled the mitochondrial genome using MitoHiFi v3.2.2 (Uliano-Silva et al. 2023) with error-corrected long-read data. We annotated it *de novo* using MitoZ v3.6 (Meng et al. 2019) and further refined the tRNA prediction with GeSeq (Tillich et al. 2017) and ARWEN (Laslett and Canbäck 2008). We removed false-positive tRNAs predicted within protein-coding sequences and combined the results from both pipelines. Finally, we visualised the mitogenome using OGDRAW (Greiner et al. 2019).

### 5. Demographic history reconstruction

We used Pairwise Sequentially Markovian Coalescent (PSMC) analysis (Li & Durbin 2011) using psmc v0.6.5-r67 to reconstruct the historical demography of *I. ampulla*. First, we generated a diploid consensus sequence by aligning short-read data to the reference genome (Assembly 6) using BCFtools (Danecek et al. 2021). We filtered variants with bcftools view-c - and converted the output to FASTQ format using vcfutils.pl, setting depth thresholds to a minimum of 6 and a maximum of 48. We then performed PSMC analysis with parameters -N25 -t15 -r5 -p “4+25*2+4+6” and with 100 bootstrap replicates.

## Results and discussion

### 1. Nuclear genome assembly statistics

We present the first genome assembly of *I. ampulla*, constructed using 21 GB of error-corrected long-read data and 118 GB of filtered short-read data. We generated initial assemblies and refined them through two rounds of QuickMerge with different assembly combinations, resulting in a total of eight assemblies (see Supplementary table 1 for detailed statistics). Among these, we selected Assembly 6 as the final assembly based on its optimal balance of contiguity and completeness. The final assembly spans 1.98 GB, consisting of 8,792 contigs with an N50 value of 632 kb. BUSCO analysis indicated 90.3% genome completeness, and we estimated the GC content to be 36.35% (Figure 1-B). The genome size of *I. ampulla* falls well within the known range for molluscs, which exhibit considerable variation, from less than 1 GB in Biomphalaria glabrata (850.6 Mb; accession no. GCA_947242115.1) and Elysia timida (754.4 Mb; accession no. GCA_043644045.1) to over 5 GB in Oreohelix idahoensis (5.5 Gb; accession no. GCA_024509875.1). This diversity highlights the broad range of genome sizes within the parent subclass Heterobranchia.

**Figure 1.**
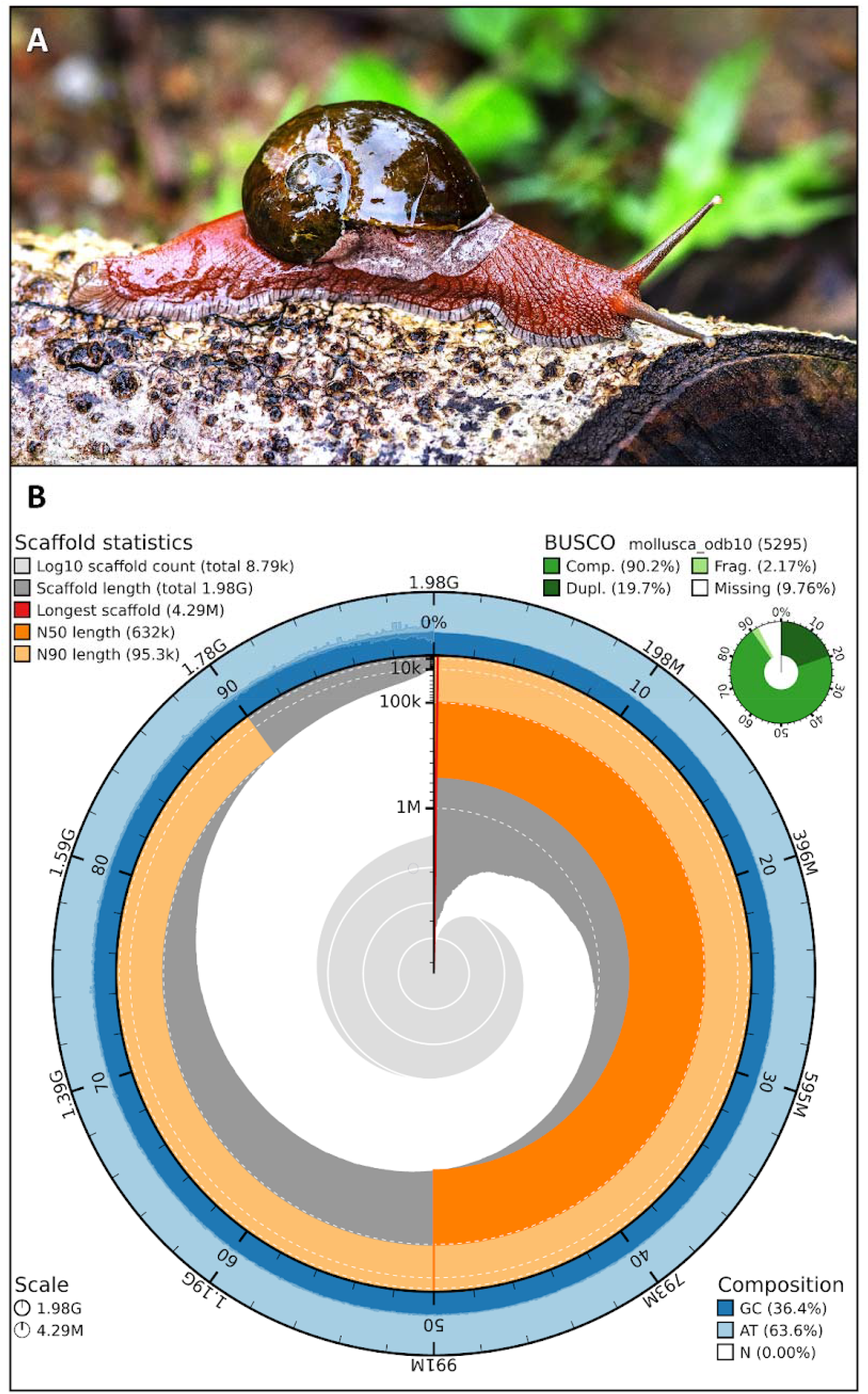
A) A red morph individual of *Indrella ampulla*. B) Snail plot of the final assembly of *Indrella ampulla* along with the assembly statistics, sequence composition, and BUSCO completeness.

### 2. Repeat masking and annotation

Repeat annotation revealed that approximately 54% of the genome consists of repeat elements. The major categories of repeats include SINEs, LINEs, LTR elements, DNA transposons, rolling circles, and simple repeats. LINEs constituted the largest proportion, accounting for 25.94% of the genome, nearly half of all repeat content (See Supplementary table 2 for complete repeat statistics). Gene prediction using GALBA identified a total of 20,418 protein-coding genes. BUSCO analysis of the predicted proteome indicated 88.7% completeness, with a duplication rate of 17%. Consistency analysis using OMArk showed that 68.03% of the proteins were consistently placed within the clade Mollusca, while 20.24% were classified as unknown (Figure 2-A). Importantly, OMArk predicted no contaminants in the genome.

**Figure 2.**
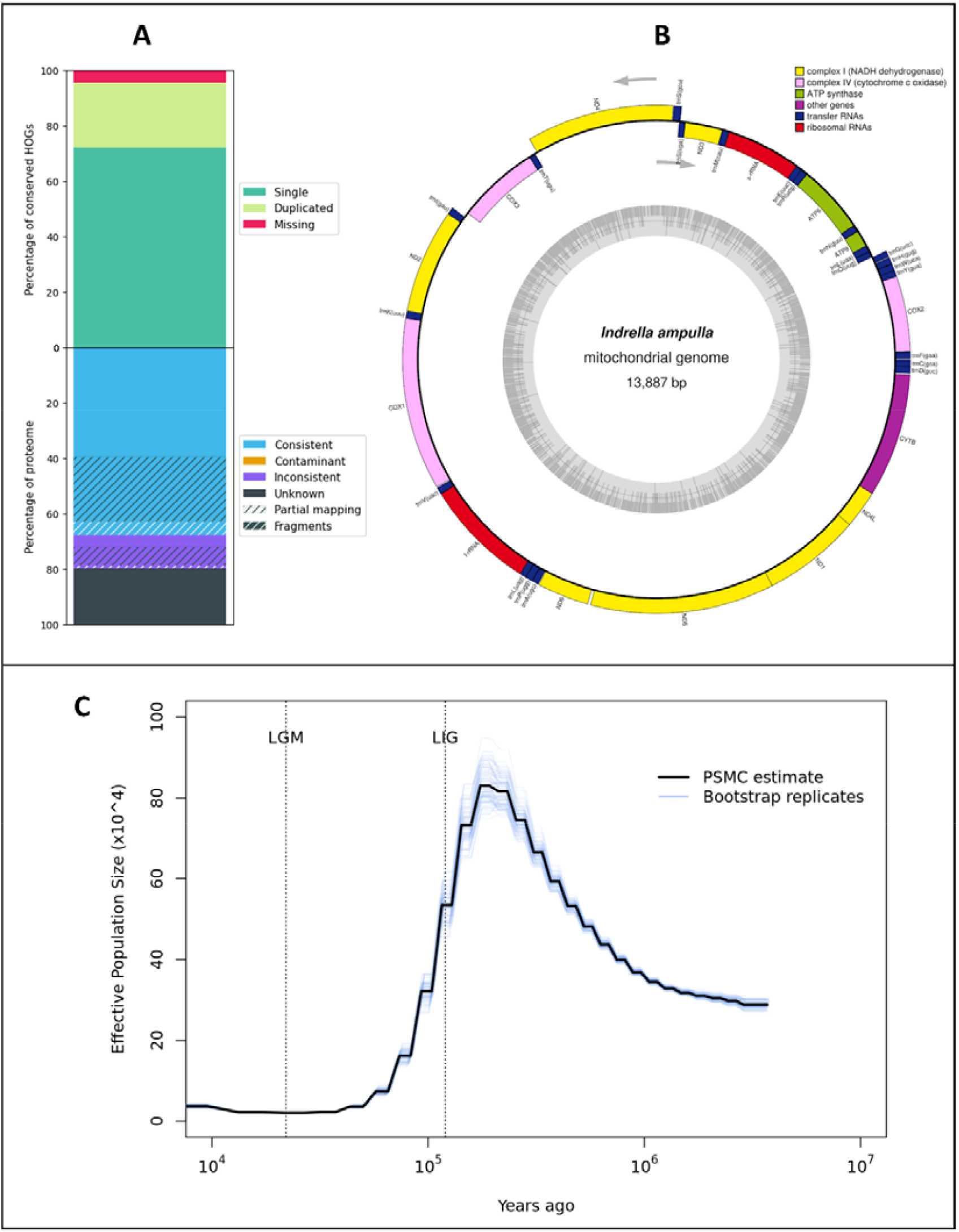
A) OMArk analysis of annotated proteome quality assessment for the *Indrella ampulla* genome. B) Circular map of the *Indrella ampulla* mitochondrial genome assembly, showing the gene arrangement, protein-coding regions, rRNAs, and tRNAs. The inner circle illustrates the GC content distribution across the mitogenome. Arrows on the outer and inner sides of the circle denote directions of gene transcription. C) PSMC plot illustrating historical effective population size trends (black line) of *Indrella ampulla* over time. The X-axis represents time in years before the present, scaled by the generation time (g) and mutation rate (μ), and the Y-axis shows the effective population size (Ne). LGM represents Last Glacial Maximum and LIG represents Last Interglacial period.

### 3. Mitogenome assembly statistics

The mitochondrial genome assembly is 13,887 bp long and is one circular contig (Figure 2-B). It contains 13 protein-coding genes, 22 tRNA genes, and two ribosomal RNA genes. Four protein-coding genes (ATP6, ATP8, ND3 and COX3) are present in the minor strand, while the rest are in the major strand. Among the tRNAs, 14 are present in the plus/major strand, while eight are present in the minor strand. The 13 protein-coding genes include 3 Cytochrome oxidase genes, 1 Cytochrome b, 7 NADH genes and 2 ATP synthase genes. The size of the mitochondrial genome of *I. ampulla* is comparable to other molluscs; for example, 13,670 bp in *Biomphalaria glabrata* (DeJonget al., 2004), 14,100 bp in *Cepaea nemoralis* (Terrett et al. 1996), and 15,057 bp in *Achatina fulica* (He et al. 2016).

### 4. Demographic history

We observed a sudden drop in Effective population size (N_e_), concurrent with the beginning of the last glacial period (Figure 2-C). The Pleistocene glacial cycles are major drivers of various intricate phylogeographic patterns across the globe. However, its role in shaping the genetic diversity of flora and fauna of the Western Ghats is unclear. Some studies suggest a change in rainfall patterns and increase in aridification before and during the last glacial maxima (Srivastava et al. 2015; Raja et al. 2018), which can potentially lead to population decline in forest-dwelling species. Several co-distributed species, such as passerine birds, exhibit signatures of population decline around the same time (Robin et al. 2010). Interestingly, in the case of the study species, the population decline began much earlier. A striking parallel has been observed in the Asian elephant populations in the Western Ghats, based on genomic data (Khan et al. 2024). Previous attempts to assess the demographic history using genetic data were ambiguous (Chakraborthy et al., 2024), underlining the importance of incorporating genomic data to assess biodiversity patterns in tropical Asia.

## Acknowledgements

The authors acknowledge the Kerala Forest Department for permission and the CCMB sequencing facility as well as Novelgene.INC. Whole genome sequencing were supported by internal funding from the National Institute of Science Education and Research (under Dept. of Atomic Energy) and SERB CRG Grant (CRG_2021_000261) awarded to ADR. NAA would like to thank the Science and Engineering Research Board (SERB), Govt. of India, for funding (EMR/2017/000199). GK was supported by a PhD fellowship from the Council of Scientific and Industrial Research, Govt. of India. The authors would also like to thank Dr. Kushankur Bhattacharyya for capturing the photograph of *I. ampulla*.

## Data Availability

The sequences and draft genome generated for this study can be found under NCBI BioProject PRJNA1227711. The short-read and long-read data are archived under SRA IDs SRR32476888 and SRR32476889, respectively. The mitogenome has been submitted to NCBI’s BankIt under the accession ID PV650439.

## Literature cited

1. Aravind NA, Rajashekhar KP, Madhyastha NA. 2005. Species diversity, endemism and distribution of land snails of the Western Ghats, India. Records of the Western Australian Museum Supplement. 68:31–38.

2. Astashyn A, et al. 2024. Rapid and sensitive detection of genome contamination at scale with FCS-GX. Genome Biol. 25:60.

3. Brůna T, et al. 2023. Galba: genome annotation with miniprot and AUGUSTUS. BMC bioinformatics. 24:327.

4. Cantalapiedra CP, Hernández-Plaza A, Letunic I, Bork P, Huerta-Cepas J. 2021. eggNOG-mapper v2: functional annotation, orthology assignments, and domain prediction at the metagenomic scale. Mol Biol Evol. 38:5825–9.

5. Chakraborty S, et al. 2024. Phylogeographical patterns are governed by geography in endemic polymorphic snail Indrella ampulla (Gastropoda: Ariophantidae). Biol J Linn Soc Lond. 142:44–57.

6. Chakraborty M, Baldwin-Brown JG, Long AD, Emerson JJ. 2016. Contiguous and accurate de novo assembly of metazoan genomes with modest long read coverage. Nucleic Acids Res. 44:e147.

7. Chakraborty S, Saha A, Aravind NA. 2020. Comparison of DNA extraction methods for non-marine molluscs: is the modified CTAB DNA extraction method more efficient than DNA extraction kits? 3 Biotech. 10:69.

8. Challis R, Richards E, Rajan J, Cochrane G, Blaxter M. 2020. BlobToolKit–interactive quality assessment of genome assemblies. G3 (Bethesda). 10:1361–74.

9. Chen S. 2023. Ultrafast one-pass FASTQ data preprocessing, quality control, and deduplication using fastp. Imeta. 2:e10

10. Cuthill IC, et al. 2017. The biology of colour. Science. 357:eaan0221.

11. Danecek P, et al. 2021. Twelve years of SAMtools and BCFtools. Gigascience. 10:giab008.

12. DeJong RJ, Emery AM, Adema CM. 2004. The mitochondrial genome of Biomphalaria glabrata (Gastropoda: Basommatophora), intermediate host of Schistosoma mansoni. J Parasitol. 90:991–997.

13. Flynn JM, et al. 2020. RepeatModeler2 for automated genomic discovery of transposable element families. Proc Natl Acad Sci. U.S.A. 117:9451-7.

14. Greiner S, Lehwark P, Bock R. 2019. OrganellarGenomeDRAW (OGDRAW) version 1.3. 1: expanded toolkit for the graphical visualization of organellar genomes. Nucleic Acids Res. 47:W59–64.

15. Gurevich A, Saveliev V, Vyahhi N, Tesler G. 2013. QUAST: quality assessment tool for genome assemblies. Bioinformatics. 29:1072–5.

16. Hayes KA, et al. 2009. Molluscan models in evolutionary biology: apple snails (Gastropoda: Ampullariidae) as a system for addressing fundamental questions. Am Malacol Bull. 27:47–58.

17. He ZP, et al. 2016. Complete mitochondrial genome of the giant African snail, Achatina fulica (Mollusca: Achatinidae): a novel location of putative control regions (CR) in the mitogenome within Pulmonate species. Mitochondrial DNA A: DNA Mapp Seq Ana. 27:1084–1085.

18. Huerta-Cepas J, et al. 2019. eggNOG 5.0: a hierarchical, functionally and phylogenetically annotated orthology resource based on 5090 organisms and 2502 viruses. Nucleic Acids Res. 47:D309–14.

19. Khan A, et al. 2024. Divergence and serial colonization shape genetic variation and define conservation units in Asian elephants. Curr Biol. 34:4692–703.

20. Kolmogorov M, Yuan J, Lin Y, Pevzner PA. 2019. Assembly of long, error-prone reads using repeat graphs. Nat Biotechnol. 37:540–6.

21. Laslett D, Canbäck B. 2008. ARWEN: a program to detect tRNA genes in metazoan mitochondrial nucleotide sequences. Bioinformatics. 24:172–5.

22. Li H, Durbin R. 2011. Inference of human population history from individual whole-genome sequences. Nature. 475:493–6.

23. Mak QC, Wick RR, Holt JM, Wang JR. 2023. Polishing de novo nanopore assemblies of bacteria and eukaryotes with FMLRC2. Mol Biol Evol. 40:msad048

24. Manni M, Berkeley MR, Seppey M, Simão FA, Zdobnov EM. 2021. BUSCO update: novel and streamlined workflows along with broader and deeper phylogenetic coverage for scoring of eukaryotic, prokaryotic, and viral genomes. Mol Biol Evol. 38:4647–54.

25. Meng G, Li Y, Yang C, Liu S. 2019. MitoZ: a toolkit for animal mitochondrial genome assembly, annotation and visualization. Nucleic Acids Res. 47:e63

26. Narayanan, S., Aravind, N. A. 2021. Observations on natural diet and reproductive behaviour of an endemic snail Indrella ampulla (Benson 1850) (Gastropoda: Ariophantidae) from the Western Ghats, India. Journal of Natural History. 55:2961–2972.

27. Nevers Y, et al. 2025. Quality assessment of gene repertoire annotations with OMArk. Nat Biotechnol. 43:124–33.

28. Ponder WF, Lindberg DR, Ponder JM. 2019. Gastropoda I: introduction and the stem groups. Biology and evolution of the Mollusca. 2:289–363.

29. Raja P, et al. 2019. Tropical rainforest dynamics and palaeoclimate implications since the late Pleistocene, Nilgiris, India. Quat Res. 91:367–82.

30. Robin VV, Sinha A, Ramakrishnan U. 2010. Ancient geographical gaps and paleoclimate shape the phylogeography of an endemic bird in the sky islands of southern India. PLoS One. 5:e13321.

31. San-Jose LM, Roulin A. 2017. Genomics of colouration in natural animal populations. Philos Trans R Soc Lond B Biol Sci. 372:20160337.

32. Smit AF, Hubley R, Green P. 2013–2015. RepeatMasker Open-4.0 [https://www.repeatmasker.org].

33. Srivastava G, et al. 2016. Monsoon variability over Peninsular India during Late Pleistocene: Signatures of vegetation shift recorded in the terrestrial archive from the corridors of Western Ghats. Palaeogeogr Palaeoclimatol Palaeoecol. 443:57–65.

34. Terrett JA, Miles S, Thomas RH. 1996. Complete DNA sequence of the mitochondrial genome of Cepaea nemoralis (Gastropoda: Pulmonata). J Mol Evol. 42:160–168.

35. Tillich M, et al. 2017. GeSeq–versatile and accurate annotation of organelle genomes. Nucleic Acids Res. 45:W6–11.

36. Uliano-Silva M, et al. 2023. MitoHiFi: a Python pipeline for mitochondrial genome assembly from PacBio high-fidelity reads. BMC bioinformatics. 24:288

37. Zimin AV, et al. 2013. The MaSuRCA genome assembler. Bioinformatics. 29:2669–77.

