## Supplementary file for "Nuclear and mitochondrial genome assemblies of *Indrella ampulla*, a terrestrial gastropod endemic to the Western Ghats"

**Author information**

Gopi Krishnan, Maitreya Sil^,^*, N. A. Aravind, Govindhaswamy Umapathy, Aniruddha Datta-Roy

**Supplementary data**

**Supplementary table 1: Overview of genome assembly statistics and quality metrics. The final assembly selected (Assembly 6) is highlighted.**

| **Assembly** | **Tool** | **Assembly type** | **No. of contigs** | **Total length (bp)** | **N50 (bp)** | **N90 (bp)** | **Largest contig (bp)** | **BUSCO stats** |
| --- | --- | --- | --- | --- | --- | --- | --- | --- |
| Assembly1 | Flye | Long read only | 19650 | 1,969,612,814 | 299,607 | 53,742 | 1,975,331 | **C:91.0%** [S:75.9%, D:15.1%], F:2.5%, M:6.5%, n:5295, E:3.2% |
| Assembly2 | Masurca | Hybrid assembly | 13978 | 1,862,620,062 | 242,354 | 62,052 | 2,095,289 | **C:87.4%** [S:74.1%, D:13.3%], F:3.6%, M:9.0%, n:5295, E:3.0% |
| Assembly3 | Quickmerge | Assembly1 + Assembly2 | 17037 | 2,021,768,191 | 501,334 | 65,994 | 3,386,671 | **C:90.6%** [S:71.7%, D:18.9%], F:1.9%, M:7.5%, n:5295, E:3.2% |
| Assembly4 | Quickmerge | Assembly2 + Assembly1 | 8574 | 1,943,183,742 | 641,371 | 95,998 | 3,862,327 | **C:89.2%** [S:70.2%, D:19.0%], F:2.1%, M:8.7%, n:5295, E:3.0% |
| Assembly5 | Quickmerge | Assembly1 + Assembly3 | 17234 | 2,058,279,009 | 501,906 | 66,191 | 4,414,938 | **C:91.4%** [S:71.8%, D:19.6%], F:1.9%, M:6.7%, n:5295, E:3.2% |
| **Assembly6** | **Quickmerge** | **Assembly2 + Assembly3** | **8792** | **1,982,480,270** | **632,150** | **95,297** | **4,290,409** | **C:90.3% [S:70.6%, D:19.7%], F:2.2%, M:7.5%, n:5295, E:3.0%** |
| Assembly7 | Quickmerge | Assembly1 + Assembly4 | 14953 | 2,072,966,220 | 639,242 | 80,829 | 3,862,327 | **C:91.7%** [S:71.0%, D:20.7%], F:1.8%, M:6.5%, n:5295, E:3.1% |
| Assembly8 | Quickmerge | Assembly2 + Assembly4 | 9054 | 2,012,425,495 | 632,486 | 92,808 | 3,862,327 | **C:90.6%** [S:69.7%, D:20.9%], F:2.2%, M:7.2%, n:5295, E:3.0% |

**Supplementary table 2: Summary of repeat content characterisation**

| **Repeat type** | **Number of elements** | **Length occupied (kbp)** | **Percentage of sequence covering the genome** |
| --- | --- | --- | --- |
| SINE | 121,938 | 20,081 | 1.01 |
| LINE | 1,338,278 | 514,314 | 25.94 |
| LTR elements | 18,306 | 5,452 | 0.28 |
| DNA transposons | 219,394 | 87,562 | 4.42 |
| Rolling-circles | 5,983 | 1,523 | 0.08 |
| Unclassified | 1,896,320 | 368,417 | 18.58 |
| Small RNA | 106,312 | 18,514 | 0.93 |
| Satellites | 5,524 | 1,533 | 0.08 |
| Simple repeats | 549,950 | 41,624 | 2.10 |
| Low complexity | 67,524 | 5,342 | 0.27 |
| **Total** | **4,329,529** | **1,044,281** | **53.69** |
